# Developmental Changes in EEG Phase Amplitude Coupling and Phase Preference over the First Three Years After Birth

**DOI:** 10.1101/818583

**Authors:** Michael G. Mariscal, April R. Levin, Laurel J. Gabard-Durnam, Helen Tager-Flusberg, Charles A. Nelson

## Abstract

The coupling of the phase of slower electrophysiological oscillations with the amplitude of faster oscillations, termed phase-amplitude coupling (PAC), is thought to facilitate dynamic connectivity in the brain. Though the brain undergoes dramatic changes in connectivity during the first few years of life, how PAC changes through this developmental period has not been studied. Here, we examined PAC through electroencephalography (EEG) data collected longitudinally during an awake, eyes-open EEG collection paradigm in 98 children between the ages of 3 months and 3 years. We implement a novel technique developed for capturing both PAC strength and phase preference (i.e., where in the slower oscillation waveform the faster oscillation shows increased amplitude) simultaneously, and employed non-parametric clustering methods to evaluate our metrics across a range of frequency pairs and electrode locations. We found that frontal and occipital PAC, primarily between the alpha-beta and gamma frequencies, increased from early infancy to early childhood (p = 1.35 x 10^-5^). Additionally, we found frontal gamma coupled with the trough of the alpha-beta waveform, while occipital gamma coupled with the peak of the alpha-beta waveform. This opposing trend may reflect each region’s specialization towards feedback or feedforward processing, respectively.

**Significance Statement:** The brain undergoes significant changes in functional connectivity during infancy and early childhood, enabling the emergence of higher-level cognition. Phase-amplitude coupling (PAC) is thought to support the functional connectivity of the brain. Here, we find PAC increases from 3 months to 3 years of age. We additionally report the frontal and occipital brain areas show opposing forms of PAC; this difference could facilitate each region’s tendency towards bottom-up or top-down processing.

## Introduction

Communication between spatially separated brain regions is a critical component of healthy brain function. Oscillatory brain activity, created by the synchronous firing of large ensembles of neurons, is suggested to facilitate this communication: low frequency oscillations may reflect long distance connections, while high frequency oscillations may reflect local connections (von Stein and Sarnthein, 2000). Cross frequency coupling, which describes the interaction between different oscillatory frequencies, may therefore serve to integrate information spatially and temporally (Canolty and Knight, 2010).

One form of cross-frequency coupling is Phase-Amplitude Coupling (PAC), in which the phase of the low frequency activity modulates the amplitude of the high frequency activity. Increasing evidence implicates PAC in a variety of important functional processes. PAC reflects strength of functional connectivity (Weaver et al., 2016), and PAC strength and location has been shown to shift with task demands (Voytek et al., 2010). Additionally, differences in PAC have been found in a variety of brain disorders, including schizophrenia, autism, attention deficit hyperactivity disorder, and Parkinson’s (Salimpour and Anderson, 2019), highlighting the importance of PAC for healthy brain functioning.

Moreover, underlying characteristics of PAC may reflect an area’s functional configuration within neural networks. Phase preference, or the phase of the low frequency waveform corresponding to the largest amplitude of the high frequency waveform (Figure 1), has been found to differ by brain area (Ninomiya et al., 2015), coupling frequencies, and task performance (Lega et al., 2016). Additionally, phase preference can vary between cortical layers: recordings from multielectrode arrays implanted in the striate cortex V1 of macaque monkeys found layers IV and VI to exhibit opposing phase preferences with respect to alpha oscillations (Bollimunta et al., 2011). This may relate to how these layers interact with the thalamus, where alpha rhythms are often thought to originate or project; layer IV tends to receive thalamic input, while layer VI tends to project back to the thalamus (Guillery and Sherman, 2002). Thus, it is possible that primary phase preference (as measured on the scalp) could reflect an underlying area’s dynamic functional configuration as predominantly either feedforward or feedback during the recording period. Though typical measures of PAC are agnostic to phase preference, these findings suggest that not just *if*, but *how* gamma interacts with lower frequency waveforms may reflect a region’s role in either the intake or modulation of information for neural processing.

**Figure 1:**
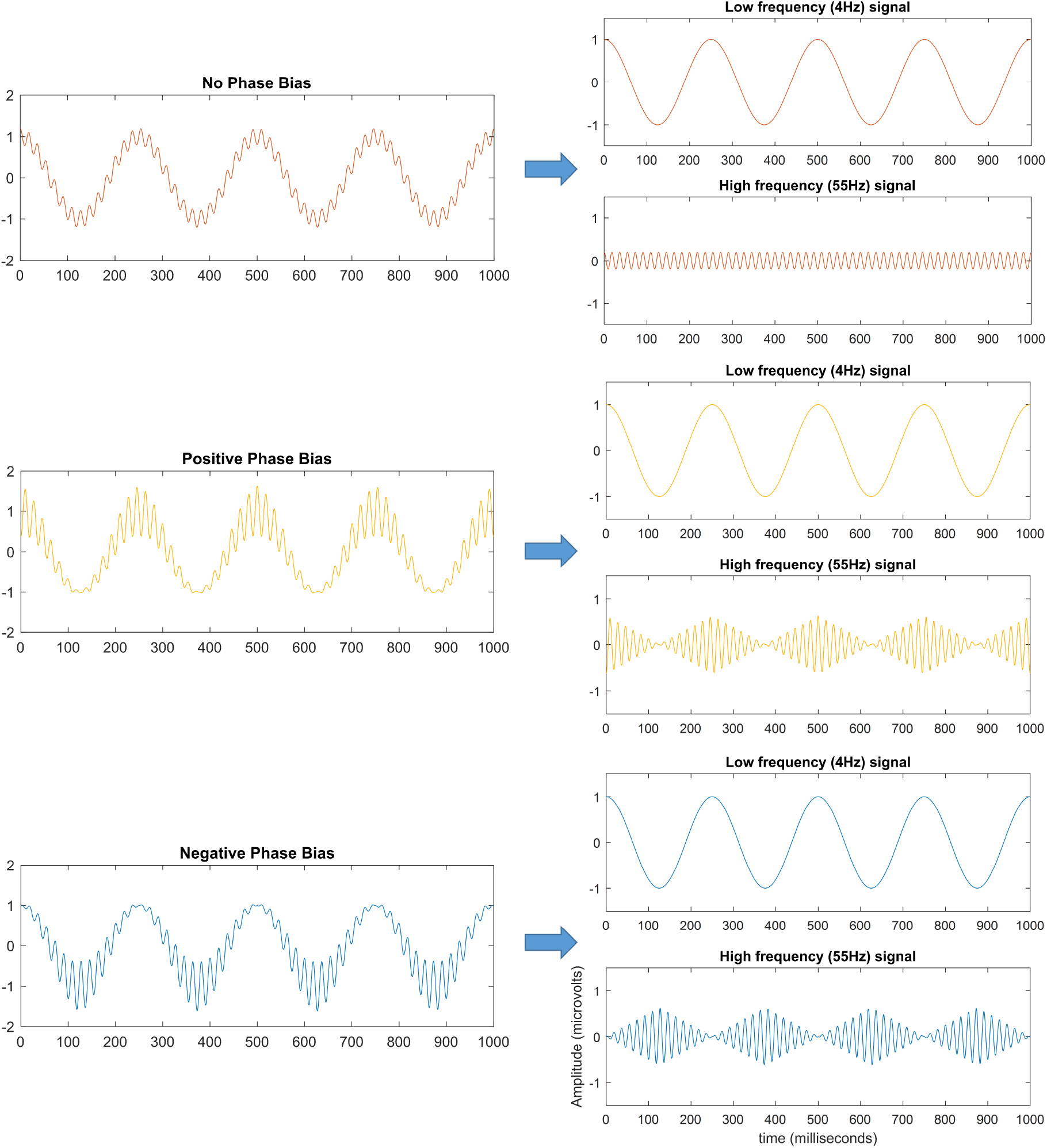
Phase-amplitude Coupling Schema. No phase bias is present when PAC is not present (red). Positive phase bias occurs when the amplitude of the high frequency signal increases at the peak of the low frequency signal waveform (yellow), and negative phase bias occurs when the amplitude of the high frequency signal increases at the trough of the low frequency signal (blue).

Dramatic developmental changes in functional connectivity have been documented across fMRI, MEG, and EEG studies (Grayson and Fair, 2017), suggesting PAC and phase preference would show developmental changes as well. However, no studies have examined the development of phase preference, and few have examined the development of PAC. Of note, theta-gamma PAC has been found to increase from 8 to 16 years in response to audio click-trains, and begins to decrease from 16 to 22 years; it has been hypothesized that this may relate to developmental changes in GABA transmission and synaptic pruning (Cho et al., 2015). Still, studies on infancy and early childhood, a period when the brain is highly plastic and shows significant structural and functional developmental changes, have not been conducted. Only one study examined PAC during infancy: PAC recorded during sleep decreased over the first two weeks after birth (Tokariev et al., 2016).

Thus, though the brain continues to demonstrate high plasticity and significant developmental changes from birth to early childhood (Gao et al., 2017), no studies have examined how PAC or phase preference develops in this age range. Here we sought to address these gaps by characterizing, through both PAC strength and phase preference, how PAC develops from infancy to early childhood. Through this process, we have also developed a novel measure reflecting both PAC strength and phase preference simultaneously.

## Methods

### Participants

The data for this study were drawn from a larger longitudinal study of neuro-cognitive development across the first 3 years of after birth. This study included later born infants who had at least one typically developing older sibling (Table 1, n = 98). The study was conducted at Boston Children’s Hospital/Harvard Medical School and Boston University. All infants had a minimum gestational age of 36 weeks, no history of prenatal or postnatal medical or neurological problems, no known genetic disorders (e.g., fragile X, tuberous sclerosis) and no family history of autism spectrum disorder or other neuropsychiatric conditions (based on parent report). Institutional review board approval was obtained from both institutions (#X06-08-0374).

**Table 1:** Participant Demographics.

|  | Participants (n=98) |
| --- | --- |
| Sex | 53(M) 45(F) |
| Maternal Education |  |
| <4 year degree | 13 (13%) |
| 4 year degree | 26 (27%) |
| Graduate degree | 49 (50%) |
| Did not answer | 10 (10%) |
| Paternal Education |  |
| <4 year degree | 19 (19%) |
| 4 year degree | 34 (35%) |
| Graduate degree | 34 (35%) |
| Did not answer | 11 (11%) |
| Race |  |
| White or Caucasian | 82 (84%) |
| Black or African American | 3 (3%) |
| Asian | 2 (2%) |
| American Indian or Alaskan Native | 0 (0%) |
| Native Hawaiian or Pacific Islander | 0 (0%) |
| More than one reported | 10 (10%) |
| Not reported | 1 (1%) |
| Ethnicity |  |
| Not Hispanic or Latino | 93 (94%) |
| Hispanic or Latino | 4 (4%) |
| Not reported | 1 (1%) |

### EEG Acquisition / Processing

Baseline EEG data were collected at 3, 6, 9, 12, 18, 24, and 36 months of age, as previously described (Gabard-Durnam et al., 2019). Infants were seated on their caregiver’s lap while a research assistant blew bubbles and/or presented toys to ensure the infant remained calm. Continuous EEG was recorded for up to 5 minutes using either a 64-channel Geodesic Sensor Net System or 128-channel Hydrocel Geodesic Sensor Nets (Electrical Geodesics, Inc., Eugene, OR). Data were sampled at either 250 or 500 Hz, and referenced at collection to a single vertex electrode (Cz). Impedances were kept below 100 kΩ (within recommended guidelines for young children, given the high-input impedance capabilities of this system’s amplifier).

Raw EEG files were exported from NetStation to MATLAB (versionR2017b, MathWorks, Natick, MA, USA) for preprocessing. Files were processed using the Batch EEG Automated Processing Platform (BEAPP) (Levin et al., 2018). Within BEAPP, the Harvard Automated Preprocessing Pipeline for EEG (HAPPE), which was developed specifically to optimize preprocessing of developmental EEG data with potentially high levels of artifact and short recordings, was used to automate preprocessing and artifact minimization (Gabard-Durnam et al., 2018). Data were first filtered using a 1 Hz high-pass filter and a 100 Hz low-pass filter. Data sampled at 500 Hz were then downsampled to 250 Hz for consistency. Only electrodes in the international 10-20 system were included in this pipeline; all other electrodes were removed before further analysis. Regions of signal with any channel’s amplitude >40 μV (the HAPPE default threshold, reflecting the reduced signal amplitude that results from wavelet-thresholding and independent components analysis in HAPPE) were removed prior to segmenting data into 2 second windows for PAC analysis. For each participant, 30 segments (60 seconds of data) were then randomly selected for further analysis; files with fewer than 30 segments of data at this stage were not analyzed (Table 2). Primary PAC metrics were then obtained using code added to the BEAPP software.

**Table 2:**
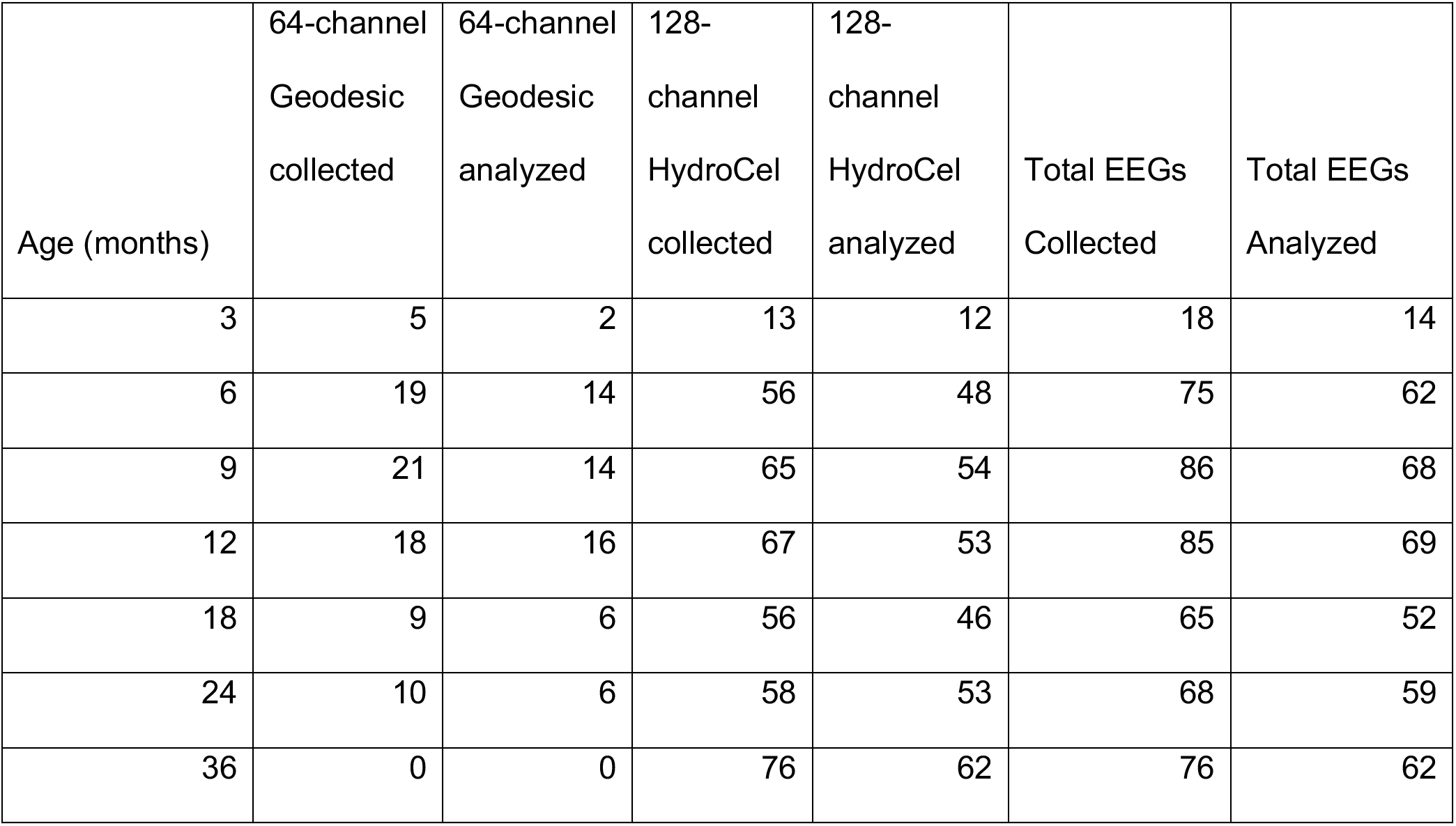
Number of EEG files (for each net type and in total) collected and analyzed per age group studied.

| Age (months) | 64-channel<br>Geodesic<br>collected | 64-channel<br>Geodesic<br>analyzed | 128-channel<br>HydroCel<br>collected | 128-channel<br>HydroCel<br>analyzed | Total EEGs<br>Collected | Total EEGs<br>Analyzed |
| --- | --- | --- | --- | --- | --- | --- |
| 3 | 5 | 2 | 13 | 12 | 18 | 14 |
| 6 | 19 | 14 | 56 | 48 | 75 | 62 |
| 9 | 21 | 14 | 65 | 54 | 86 | 68 |
| 12 | 18 | 16 | 67 | 53 | 85 | 69 |
| 18 | 9 | 6 | 56 | 46 | 65 | 52 |
| 24 | 10 | 6 | 58 | 53 | 68 | 59 |
| 36 | 0 | 0 | 76 | 62 | 76 | 62 |

Five EEG recordings were more than 3 standard deviations from the mean on one of the following HAPPE data quality output parameters: percent good channels, mean retained artifact probability, median retained artifact probability, percent of independent components rejected, and percent variance retained after artifact removal. These EEGs were evaluated for differences in overall PAC (averaged across frequencies and channels); all files were found to be within 2 SD’s of the mean at the time point of collection, so these files were included in later analyses.

### Computation of PAC Metrics

#### Modulation Index

PAC was first quantified using the Modulation Index (MI) (Tort et al., 2010). For each frequency pair, the raw signal in each segment was exported from MATLAB into Python and filtered into a low frequency and high frequency signal using code adapted from (Dupré la Tour et al., 2017). Because the frequency pairs at which PAC occurs in our age group have not been previously well defined, we began by examining PAC across a range of frequencies. To do so, for each EEG the raw signal was filtered across a range of low frequency (2-20 Hz in 2 Hz steps) and high frequency (20-100 Hz in 4 Hz steps) combinations. Subsequently, the time series of phases of the low frequency signal and the amplitude of the high frequency signal were computed. The phases of the low frequency signal were then binned into 18 20° intervals (−180° to 180°), and the mean of the amplitude of the high frequency signal occurring within each phase bin was calculated. Data were then imported into MATLAB, where the amplitude of the high frequency signal (HF_amp_) at each phase of the low frequency signal (LFϕ) was then averaged together across segments before computing the MI_raw_ as the Kullback-Leibler divergence from a uniform amplitude distribution (Tort et al., 2010). To control for factors not of interest that have been shown to affect PAC (such as spectral power), for each participant, 200 surrogate MI values (MI_surr_) were generated by repeating the procedure after offsetting HF_amp_ from LFϕ by a randomized time shift between 0.1 to 1.9 seconds. From this distribution, the mean (μ(MI_surr_)) and standard deviation were then calculated. A normalized MI (MI_norm_) was then computed as the z-score of the MI_raw_ compared to the distribution of MI_surr_ values (Canolty et al., 2007). Consequently, for each EEG, a single MI_raw_, μ(MI_surr_), and MI_norm_ were obtained.

#### Phase Bias

The Tort method used here to quantify PAC captures the presence of coupling, irrespective of where in the LFϕ the high frequency signal demonstrates increased amplitude. However, PAC can result from an increase high frequency amplitude anywhere in the low frequency waveform, including at either the positive or negative phases, corresponding to the peak and trough of the waveform respectively (Lega et al., 2016) (Figure 1). Therefore, we created a new metric to quantify the phase range (positive phases or negative phases) of the low frequency signal where the high frequency signal showed increased amplitude.

To do so, we quantified the bias of the HF_amp_ to the positive phases of LFϕ. Specifically, we defined the phase bias (ϕ bias) of a given LFϕ, HF_amp_ pair (i.e., proportion of HF_amp_ occurring in the positive phases of the LFϕ) as: ϕ bias = (ΣHF_amp_ in positive phases of the LFϕ) / Σ(HF_amp_ in all phases of the LFϕ). Thus, a value below .5 indicates the HF_amp_ shows a preference for the negative phases of LFϕ, while a value above .5 indicates the HF_amp_ shows a preference for the positive phases of LFϕ. Additionally, a larger distance from .5 (where HF_amp_ shows no preference for either positive or negative phases of LFϕ) indicates stronger ϕ bias. The ϕ bias metric was computed on the original signal to obtain the ϕ bias_raw_. Then, in the same manner MI_surr_ values were computed, ϕ bias was additionally computed after offsetting HF_amp_ from LFϕ by a randomized time shift to obtain 200 surrogate ϕ bias values (ϕ bias_surr_), of which the mean was taken (μ(ϕ bias_surr_)).

### Statistical Analysis

#### Cluster Correction

For every electrode, we now had obtained PAC values for each frequency pair. To identify statistically significant regions of interest both across frequency pairs and across channels, we first employed a clustering method. We used the clustering techniques described below to identify regions of interest for three PAC metrics. First, we clustered on MI to identify regions that exhibited PAC (PAC+). Second, we clustered on ϕ bias to identify regions where HF_amp_ is significantly increased in the negative phases of LFϕ (-LFϕ preference); i.e., ϕ bias < 0.5. Third, we clustered again on ϕ bias, this time to identify regions where HF_amp_ is increased in the positive phases of LFϕ (+LFϕ preference); i.e., ϕ bias > 0.5. All three of these techniques used data from all useable EEGs, at every age (3 months to 3 years) at which they were collected. We then modified the clustering metrics to identify regions where these three PAC metrics changed with age.

Whereas most clustering procedures involve comparing one condition to another (i.e., task vs. baseline stimuli), our task analyzed resting EEG data and therefore had no analogous comparison condition (or baseline) against which to test. In its place, we used the mean of the metric across the surrogate values (μ(MI_surr_) or μ(ϕ bias_surr_)). These data serve as an effective ‘control’ set as it retains the characteristics of the signal (power, noise), while any coupling present in the actual signal should not be retained. Thus, we used clustering against μ(MI_surr_) or μ(ϕ bias_surr_) to reveal regions where the PAC metric of interest is present to a significant degree.

#### Identification of PAC+ Regions across all ages

For our clustering procedure, we implemented a method that closely followed that by (Maris and Oostenveld, 2007).

1. For every channel, between every filtered low frequency signal and high frequency signal, a t-test was used to compare the MI_raw_ values across all files and the μ(MI_surr_) values across all files.
2. Data points where the null hypothesis was rejected (p < .05, 2-sided t-test) were selected.
3. Selected data points on the same channel adjacent to one another in terms of low frequency or high frequency were grouped together into clusters (MATLAB function bwlabel, connectivity = 4).
4. After subtracting the minimum t value needed to achieve p < .05 from all data points, cluster level statistics were computed by taking the sum of t values within each cluster.

Then, to compute which clusters were significant (i.e. unlikely to occur with that strength or size by chance, or to remove clusters that may be spurious), we implemented the following:

1. For half of the participants, selected randomly, the MI_raw_ and μ(MI_surr_) data were ‘flipped’, such that their μ(MI_surr_) data were treated as MI_raw_, and their MI_raw_ were treated as μ(MI_surr_).
2. Test statistics and cluster sizes were calculated in the same manner as used previously.
3. The previous steps were repeated 200 times, creating a distribution of cluster sizes.
4. Clusters < 95^th^ percentile of this distribution were removed from further analysis.

#### Identification of -LFϕ preference and +LFϕ preference regions across all ages

This procedure was then repeated using ϕ bias_raw_ and μ(ϕ bias_surr_) as the PAC metric, with the goal of identifying -LFϕ preference regions and +LFϕ preference regions. Unlike comparing MI_raw_ and MI_surr_, where PAC should lead to MI_raw_ being greater, not less, than MI_surr_, PAC can lead to ϕ bias_raw_ being greater than or less than μ(ϕ bias_surr_). When HF_amp_ is increased in the negative phases of LFϕ, ϕ bias_raw_ will be less than μ(ϕ bias_surr_), and when HF_amp_ is increased in the positive phases of LFϕ, ϕ bias_raw_ will be greater than μ(ϕ bias_surr_). As a result, clusters where the t-statistic is significantly negative highlight regions where HF_amp_ is increased in the negative phases of LFϕ, while clusters where the t-statistic is significantly positive highlight regions where HF_amp_ is increased in the positive phases of LFϕ. For this reason, the clustering procedure was implemented twice, once with a left-tailed t-test, and once with a right-tailed t-test (p < .025 to reject H_o_). This prevented grouping of regions where some data points exhibited -LFϕ preference and some exhibited a +LFϕ preference.

#### Development of PAC Metrics with age

Several different regions of interest survived cluster correction, where PAC metrics (MI or ϕ bias) were found to be significant when EEGs from all ages (3 months to 3 years) were included. To investigate how our PAC metrics in these regions changed with age, age was correlated with MI_norm_ averaged in each area (PAC+, -LFϕ preference, +LFϕ preference), as well as in all frequencies and channels tested. This was repeated with ϕ bias as the metric; however, because averaging ϕ bias in regions that exhibit opposing phase preference will cancel out these phase preferences, ϕ bias was only analyzed in regions grouped by phase preference (-LFϕ preference, +LFϕ preference).

The above-mentioned clustering techniques determined regions in which PAC metrics were significant when EEGs from all ages were included. However, it is possible that a different set of regions would be significant when clustering techniques were altered to explicitly determine regions in which PAC metrics demonstrated significant change across age. Therefore, we developed two additional clustering techniques that aim to capture regions where PAC metrics change with age.

#### Identification of regions where MI changes with age

We designed a clustering metric to capture regions where MI values change with age. To do so, we ran the clustering algorithm as described previously with the exception of the test statistic. For each channel and frequency pair, the MI_raw_ and μ(MI_surr_) data of each participant were collected. Data were given a label for an additional variable ‘raw’, such that MI_raw_ data were labeled as 1 and μ(MI_surr_) data as 0. A linear regression was then run, where MI was predicted by the variables age, raw, and the interaction between age and raw. The test-statistic (t value and p value) was then defined as the respective values for the coefficient of this interaction term. Consequently, a positive and significant t value indicates MI_raw_ increased more with age than μ(MI_surr_), and positive clusters therefore indicate regions where PAC is increasing with age.

#### Identification of regions where ϕ bias changes with age

This procedure was then repeated using ϕ bias_raw_ and μ(ϕ bias_surr_) as the PAC metric, with the goal of identifying regions where ϕ bias changed with age. As with clustering on ϕ bias over all ages, the clustering procedure was implemented twice, once with a left-tailed t-test, and once with a right-tailed t-test (p < .025 to reject H_o_ in each case).

## Results

### Identification of PAC+ Regions across all ages

Clustering on MI of all participants regardless of age selected 67.12% of low frequency, high frequency, and channel combinations (Figure 2, Table S1). All channels analyzed contained at least one significant cluster of PAC, with the largest clusters centered primarily occipitally (channels O1, O2, and P7) and frontally (Fp1 and Fp2).

**Figure 2:**
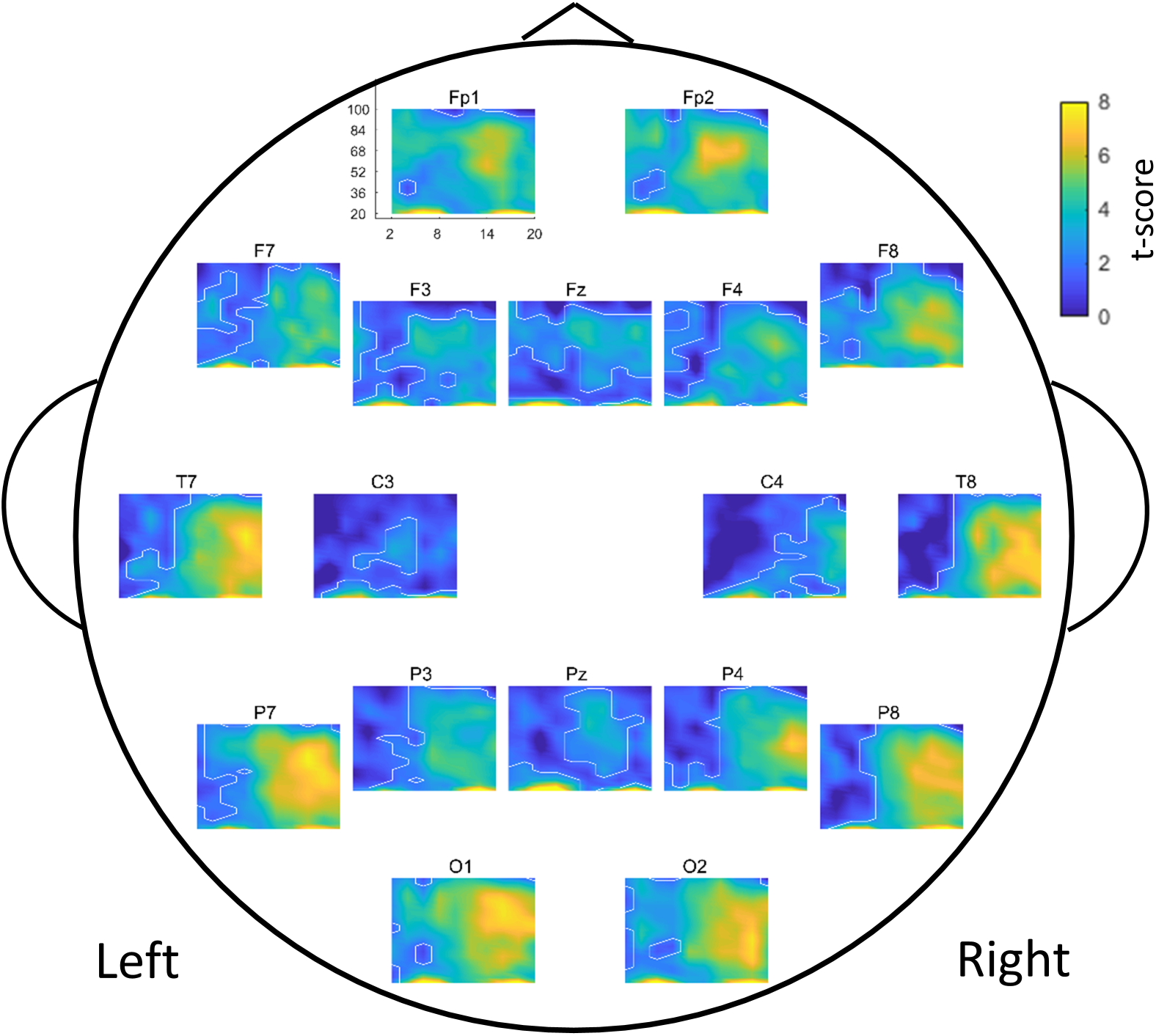
PAC+ regions across all ages. Comodulograms of t-scores, for each electrode, showing MI of each area. White lines outline regions with significant MI (PAC+ regions, p < .05, corrected for multiple comparisons). In some cases (e.g. Fp1, Fp2, O1, and O2) PAC+ regions cover most of the comodulogram; therefore, in these cases, white lines outlining blue regions mark the border of a small area that is not PAC+. Comodulograms indicate the level of coupling between phase frequencies (x-axis, 0-20Hz), and amplitude frequencies (y-axis, 20-100Hz). Each electrode is plotted at approximate electrode location on scalp. All analyzed EEGs were included (regardless of age at collection).

### Identification of -LFϕ preference and +LFϕ preference regions across all ages

Clustering on ϕ bias of all participants regardless of age revealed two sets of clusters: -LFϕ preference regions, which occurred primarily frontally, where ϕ bias was significantly less than μ(ϕ bias_surr_), and +LFϕ preference regions, which occurred more posteriorly, where ϕ bias was significantly greater than μ(ϕ bias_surr_) (Figure 3a, Table S2). Phase amplitude plots for each area confirm each area shows opposing phase preference. -LFϕ preference regions demonstrate a peak in high frequency amplitude occuring at approximately −90° low frequency phase, as well as a trough occurring at +90° (Figure 3b), while +LFϕ preference regions show a peak in high frequency amplitude occurring at approximately +90° low frequency phase, and a trough occurring at −90° phase (Figure 3c).

**Figure 3:**
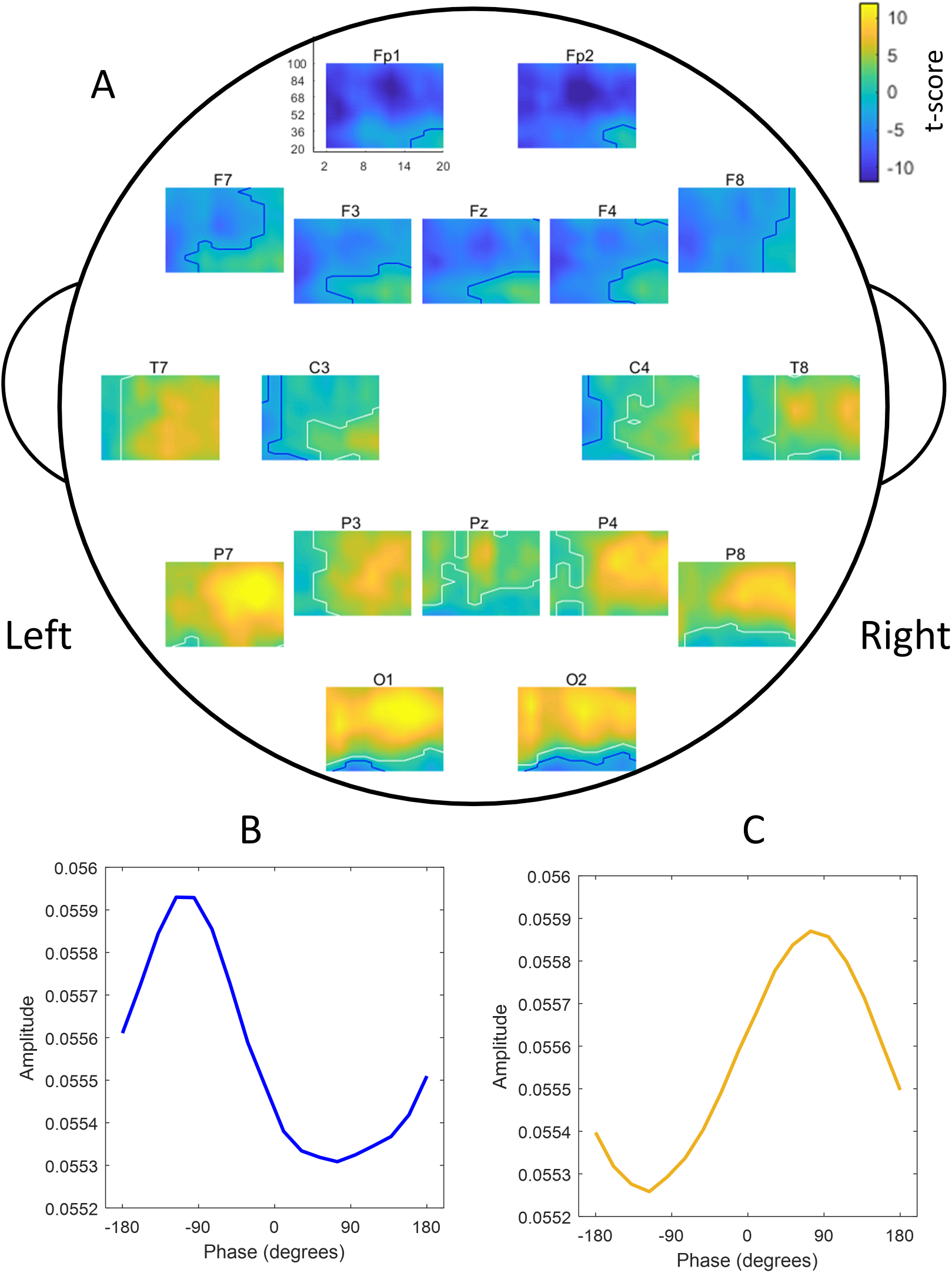
-LFϕ preference and +LFϕ preference regions across all ages, and resulting phase-amplitude plots. **A)** Comodulograms of t-scores, for each electrode, showing ϕ bias of each area. White lines outline regions where ϕ bias is high (+LFϕ preference regions, yellow) while blue lines outline regions where ϕ bias is low (-LFϕ preference regions, blue) (p < .05, corrected for multiple comparisons). All participants were included (regardless of age). **B)** Amplitude as a function of phase, averaged over all frequency pairs and channels belonging to clusters where ϕ bias is negative (left) and positive (right). All participants were included (regardless of age).

### Development of PAC Metrics with age

We then investigated how our PAC metrics in these clusters changed with age (Table 3). MI_norm_ in PAC+ regions increases with age (Table 3, Figure 4) and does not differ as a function of net type (p > .1). Changes in ϕ bias depend on region (determined regardless of age): in -LFϕ preference regions, ϕ bias decreases with age, while in +LFϕ preference regions, ϕ bias increases with age (Table 3, Figure 5). ϕ bias in these two regions has a strong inverse correlation (r = 0.612, p = 4.25 x 10^-41^).

**Figure 4:**
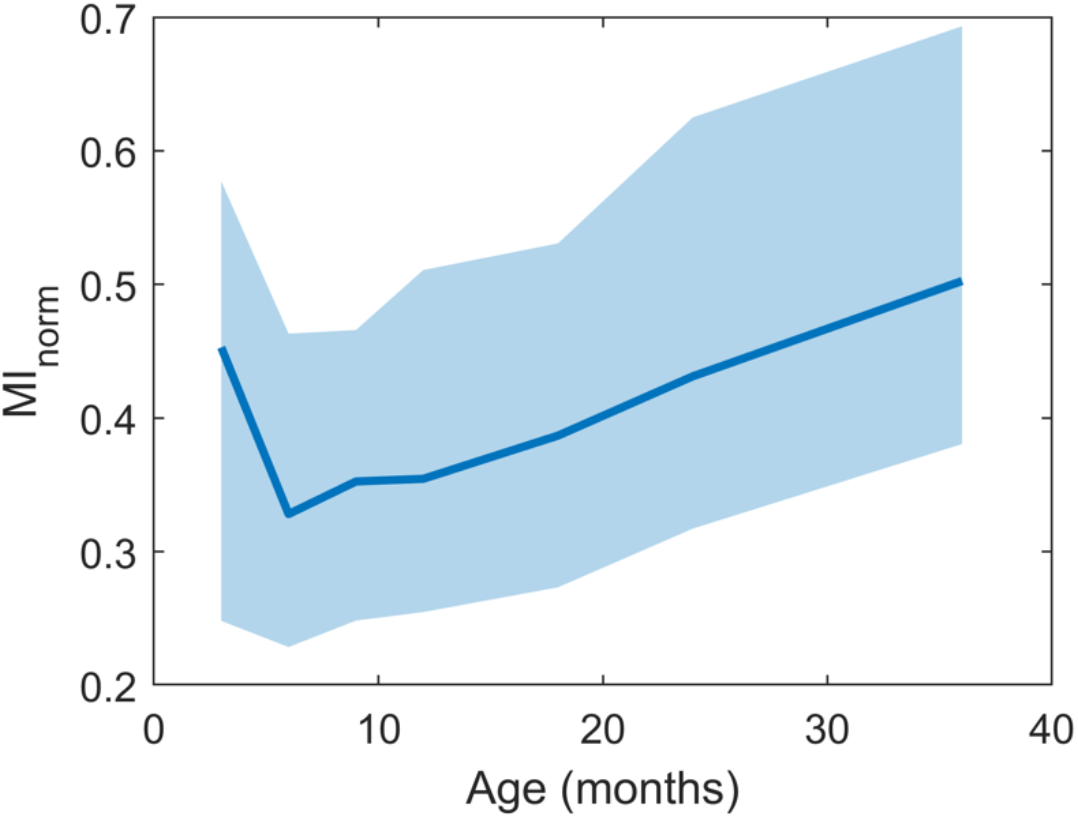
Development of MI_norm_ with age. Median MI_norm_ in PAC+ regions plotted as a function of age. Regions between the 25^th^ and 75^th^ percentiles are shaded.

**Figure 5:**
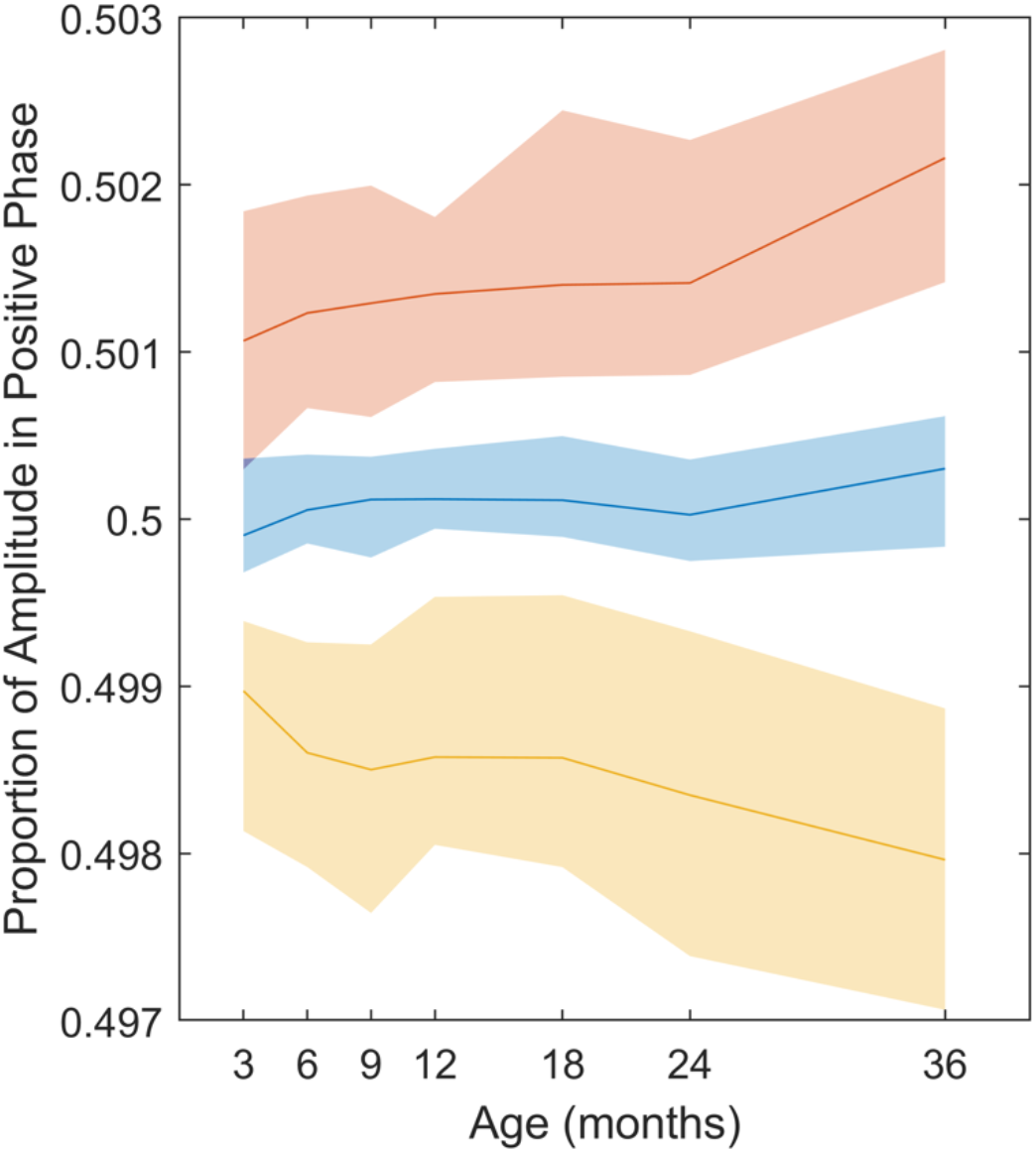
Development of ϕ bias with age. ϕ bias is plotted as a function of age, in all regions (blue), -LFϕ preference regions (yellow), and +LFϕ preference regions (red).

**Table 3:** Correlations between PAC metrics (averaged across specific regions) and age.

| Metric | Area | R | P Value |
| --- | --- | --- | --- |
| MI<sub>norm</sub> | All | 0.213 | 2.69x10<sup>-5</sup> |
| MI<sub>norm</sub> | PAC+ | 0.220 | 1.35x10<sup>-5</sup> |
| MI<sub>norm</sub> | -LFϕ preference | 0.136 | 0.00759 |
| MI<sub>norm</sub> | +LFϕ preference | 0.221 | 1.25x10<sup>-5</sup> |
| $\phi$ bias | -LFϕ preference | -0.158 | 0.0018 |
| $\phi$ bias | +LFϕ preference | 0.270 | 7.77x10<sup>-8</sup> |

### Identification of regions where MI changes with age

We then conducted an analysis of all electrodes and frequency pairs to determine which regions showed significant changes in MI with age. All regions found to be significant in this analysis showed an increase (as opposed to a decrease) in MI with age. We found increases in MI occurred most prominently in the posterior regions (Figure 6, Table S3). Specifically, we found the largest cluster in the alpha-beta and gamma frequency pairs centered on O2, and extending to electrodes O1, Pz, P8, and P7.

**Figure 6:**
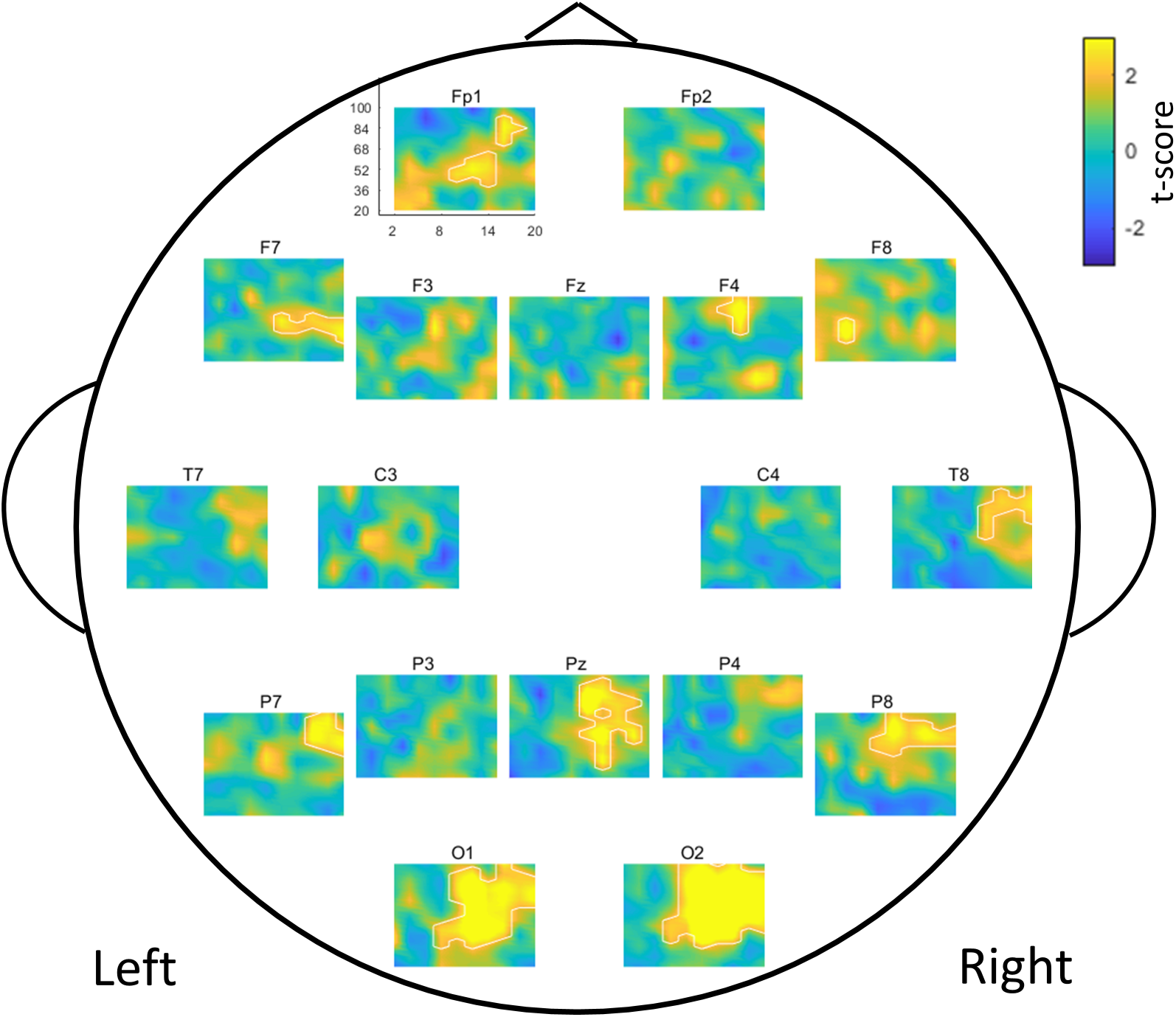
Regions where MI changes with age. Comodulograms of t-scores, for each electrode, showing regions where MI changes with age. White lines outline regions that increase with age (p < .05, corrected for multiple comparisons).

### Identification of regions where ϕ bias changes with age

We then examined which regions showed changes in ϕ bias with age. In posterior regions, ϕ bias showed similar increases with age as MI (Figure 7). Additionally, ϕ bias showed a group of negative clusters largely anteriorly, highlighting regions which develop a stronger preference for the negative phases of LFϕ with age (Table S4).

**Figure 7:**
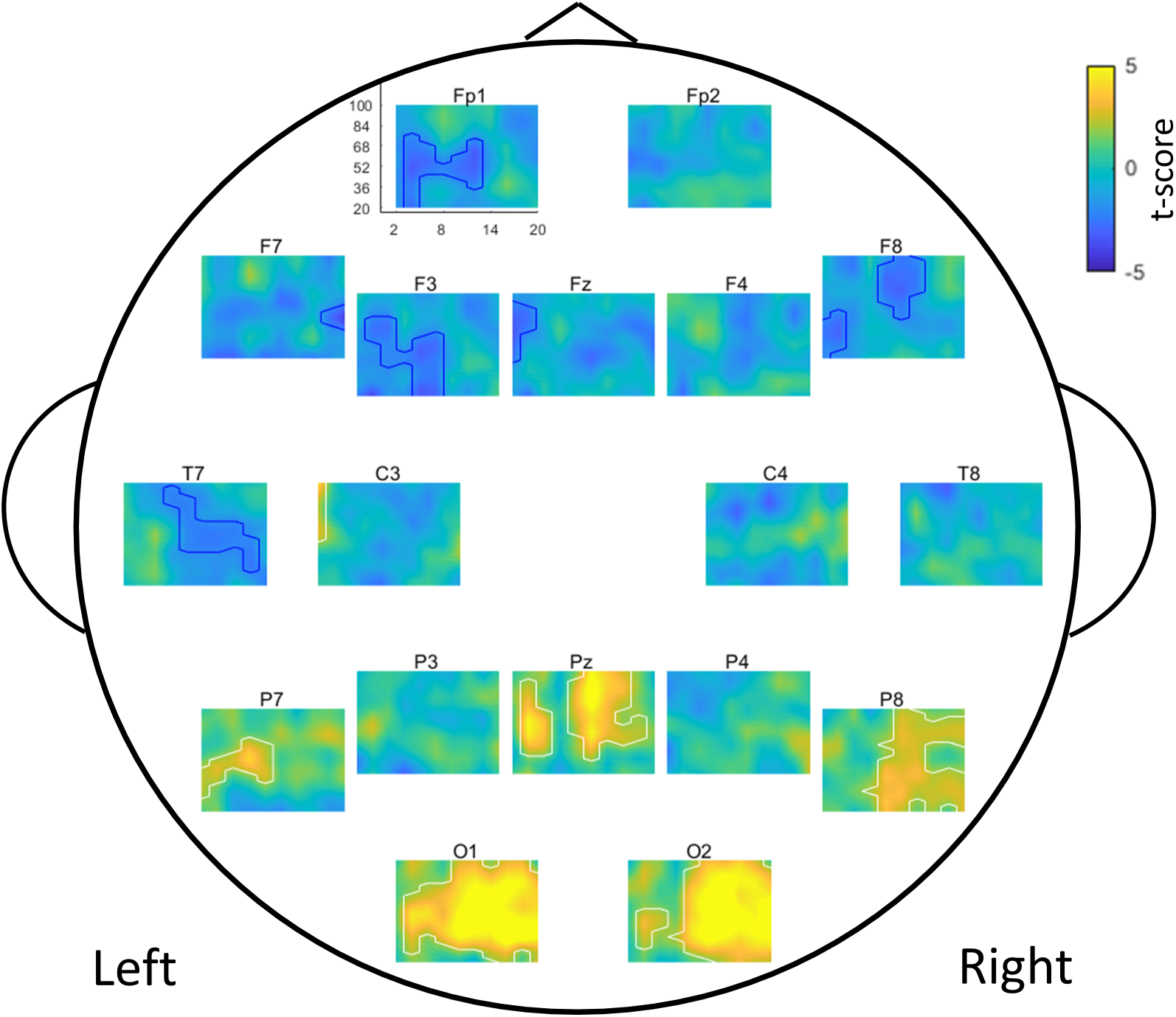
Regions where ϕ bias changes with age. Comodulograms of t-scores, for each electrode, showing regions where ϕ bias changes with age. White lines outline regions where ϕ bias increased with age, and blue lines outline regions where ϕ bias decreased with age (p < .05, corrected for multiple comparisons).

## Discussion

Though the brain exhibits significant changes in functional connectivity during infancy and early childhood, how PAC and its underlying phase preference develops over this period has not been studied. Here, we demonstrate the typical development of PAC from 3 months through 3 years of age. This PAC occurs most prominently between alpha-beta and gamma, largely consistent with several reports of alpha-gamma PAC in resting state recordings (Roux et al., 2013; Berman et al., 2015; Gohel et al., 2016), but is also present between theta and beta, theta and gamma, and alpha and beta. We observe PAC broadly across the scalp, suggesting a relatively ubiquitous presence of cross-frequency coupling. The phase preference of this PAC shows opposing trends, separated by scalp region: PAC in posterior areas (particularly over occipital lobes) is driven by a peak in gamma amplitude during the positive phases of LFϕ, while PAC in anterior areas (particularly over frontal lobes) is driven by a peak in gamma amplitude during the negative phases of the LFϕ. Though these areas appear to demonstrate similar PAC as indicated by MI, these two forms can be separated effectively using a measure of ϕ bias newly described above.

It is possible that phase bias may reflect the functional specialization of feedback (e.g., corticothalamic) and feedforward (e.g., thalamocortical) activity in the frontal and occipital areas respectively. Gamma amplitude is maximal at the peak (90°) of the alpha-beta wave in the occipital cortex, but gamma amplitude is maximal at the trough (−90°) of the alpha-beta wave in the frontal cortex. It has been previously described that cortical layer 4 (which predominantly accepts thalamic input) and layer 6 (which predominantly provides thalamic input) (Guillery and Sherman, 2002) exhibit opposing phase preferences in respect to alpha oscillations (Bollimunta et al., 2011). This functional specialization is also reflected structurally: layer IV tends to be thick posteriorly but thin anteriorly in the cortex, whereas layer VI tends to be thick in frontal cortex but thin in occipital and other predominantly sensory cortices (Wagstyl et al., 2019). Scalp recordings cannot classically differentiate among the cortical layers that may be driving findings in particular electrodes. In the context of these prior findings, however, the possibility should be considered that gamma amplitude synchronizing with the alpha-beta peak (i.e., +LFϕ preference) could reflect a predominance of feedforward, bottom-up processes (e.g., processing of visual information entering the occipital cortex), while synchronization with the alpha-beta trough could reflect a predominance of feedback, top-down processes (e.g., internal regulation of attention by the frontal cortex). In this case, the findings would thus suggest that over the first years of life, there is a specialization of regions for their particular functions: while there is indeed a developmental shift towards primarily feedforward activity with age measured over occipital regions (where incoming visual information is processed) there is also a shift towards more feedback activity measured over frontal regions (which provide top-down regulation of how the brain processes information).

An alternative (though not necessarily incongruous) explanation for the opposing phase preferences across regions involves a dipole effect. Dipole effects are frequently observed in EEG data (Ebersole, 1991; Ochi et al., 2001) and reflect a single tangentially oriented source (Osipova et al., 2008). However, the broad (rather than more localized) dipole effect seen here would likely reflect a deep source (e.g., originating from thalamus rather than cortex). While alpha and surrounding frequencies can indeed originate in the thalamus (or involve thalamocortical circuitry) (Bollimunta et al., 2011; Seeber et al., 2019), higher-frequency activity such as gamma is typically thought to originate more locally (Buzsáki and Wang, 2012). Therefore, while a single deep source may determine or coordinate LFϕ across multiple regions, this source is unlikely to be generating the coupled HF_amp_ portion of the signal. Additionally, while high frequency activity can occur as a result of muscle artifact, we would not expect such artifactual activity to be coupled the phase of the lower frequency activity.

It has been suggested PAC may serve to connect spatially and functionally segregated regions depending on the task; here, participants were not given an explicit task. Interestingly, the electrodes and frequencies where we observe the strongest PAC largely overlap with the electrodes (O1/O2, P3/P4, P7/P8, F3/F4, and Fz) frequency bands (alpha and beta) implicated in the DMN, a network thought to be active during resting state (Buckner et al., 2008). However, due to the limited spatial resolution of EEG, alternative methods are needed to better test this hypothesis.

This study has several limitations. First, our sample size at the 3 month time point was relatively small (n=14); more participants would allow for a more exact description of PAC at this age. Additionally, because most participants did not provide useable data for all time points, the study did not analyze trajectories of PAC on an individual level. Finally, though EEG’s temporal resolution lends itself to analyzing PAC, weak spatial resolution prevents more exact description of the PAC development across brain regions and layers; research using alternative neuroimaging methods, or source analysis, could answer some of the questions we leave open here.

In summary, this study documents the emergence of PAC in early childhood. We describe a new metric for measuring phase bias, which reflects more fine-grained information about PAC than the modulation index. We find HF_amp_ is increased at opposing phases of LFϕ in anterior areas as compared to posterior areas. Future studies would be beneficial in further assessing the potential functional relevance of this metric; we suggest laminar differences in the direction of information flow (feedforward vs. feedback) as one potential avenue for further exploration.

## Acknowledgements

The present research was supported by the Child Neurology Foundation (to ARL), University of Tokyo International Research Center for Neurointelligence (to LGD); and the National Institute on Deafness and Communication Disorders (R01-DC010290 to HTF and CAN; R21 DC 08637 to HTF). We thank all the laboratory members who contributed to this study, and the participants and their families who make this research possible.

## Code Accessibility

The code to process EEG data is publically available under the BEAPP and HAPPE software licenses (BEAPP: https://github.com/lcnbeapp/beapp; HAPPE: https://github.com/lcnhappe/happe). Additional code used for calculation of metrics and statistical analysis is available upon request.

---

Glossary
| Term | Definition |
| --- | --- |
| Signals |  |
| $LF\phi$ | The phase distribution of the low frequency signal |
| $HF_{amp}$ | The amplitude distribution of the high frequency signal |
| Metrics |  |
| MI | A measure of PAC strength |
| $MI_{raw}$ | MI of the actual $HF_{amp}$ and $LF\phi$ distributions |
| $MI_{surr}$ | MI of $HF_{amp}$ offset from $LF\phi$ by a randomized time shift |
| $\mu(MI_{surr})$ | Mean of 200 $MI_{surr}$ values |
| $MI_{norm}$ | Normalized Modulation Index |
| $\phi$ bias | The proportion of $HF_{amp}$ occurring in the positive phases of the $LF\phi$ |
| $\phi$ bias<sub>raw</sub> | $\phi$ bias of the actual $HF_{amp}$ and $LF\phi$ distributions |
| $\phi$ bias<sub>surr</sub> | $\phi$ bias of $HF_{amp}$ offset from $LF\phi$ by a randomized time shift |
| $\mu(\phi$ bias<sub>surr</sub>) | Mean of 200 $\phi$ bias<sub>surr</sub> values |
| Regions |  |
| PAC+ | Regions that show significant MI across age |
| -$LF\phi$ preference | Regions where $HF_{amp}$ is significantly increased during the negative phases of the $LF\phi$ across age |
| +$LF\phi$ preference | Regions where $HF_{amp}$ is significantly increased during the positive phases of the $LF\phi$ across age |

**Table S1:** PAC+ regions. Regions demonstrating significant MI across age. Low frequency and high frequency values refer to the frequency pair demonstrating the absolute maximum t test value.

| Cluster size | Low Frequency (Hz) | High Frequency (Hz) | Channel |
| --- | --- | --- | --- |
| 50.29 | 6 | 20 | Fz |
| 52.01 | 4 | 68 | F7 |
| 57.01 | 12 | 56 | C3 |
| 59.36 | 2 | 72 | F3 |
| 62.35 | 2 | 56 | F4 |
| 83.73 | 18 | 20 | C3 |
| 91.32 | 8 | 20 | Pz |
| 150.2 | 12 | 72 | Pz |
| 280.8 | 18 | 20 | C4 |
| 317.0 | 18 | 76 | Fz |
| 392.2 | 18 | 20 | F3 |
| 466.9 | 6 | 20 | F7 |
| 472.2 | 18 | 20 | P3 |
| 472.2 | 18 | 20 | F4 |
| 559.3 | 18 | 20 | P4 |
| 636.9 | 18 | 20 | F8 |
| 678.0 | 18 | 20 | P8 |
| 678.6 | 18 | 20 | T8 |
| 689.4 | 16 | 20 | T7 |
| 801.0 | 6 | 20 | Fp1 |
| 827.1 | 18 | 20 | Fp2 |
| 842.7 | 16 | 72 | P7 |
| 844.4 | 6 | 20 | O2 |
| 943.8 | 6 | 20 | O1 |

**Table S2:** -LFϕ preference and +LFϕ preference regions. Regions demonstrating significant ϕ bias across age. -LFϕ preference regions are indicated by negative cluster sizes, while +LFϕ preference regions are indicated by positive cluster sizes. Low frequency and high frequency values refer to the frequency pair demonstrating the absolute maximum t test value.

| Cluster size | Low Frequency (Hz) | High Frequency (Hz) | Channel |
| --- | --- | --- | --- |
| -1580 | 16 | 40 | Fp2 |
| -1468 | 16 | 32 | Fp1 |
| -941.6 | 10 | 24 | Fz |
| -841.4 | 12 | 20 | F4 |
| -758.5 | 18 | 72 | F8 |
| -740.8 | 20 | 56 | F3 |
| -521.9 | 6 | 28 | F7 |
| -121.2 | 4 | 40 | C3 |
| -111.0 | 4 | 76 | C4 |
| -103.5 | 20 | 20 | O2 |
| -32.13 | 8 | 28 | O1 |
| 195.2 | 20 | 36 | C3 |
| 432.9 | 20 | 44 | C4 |
| 475.3 | 12 | 80 | Pz |
| 639.4 | 18 | 68 | T8 |
| 888.5 | 14 | 56 | P3 |
| 895.2 | 12 | 28 | T7 |
| 1014 | 18 | 72 | P8 |
| 1056 | 12 | 72 | P4 |
| 1207 | 12 | 76 | O2 |
| 1375 | 14 | 80 | O1 |
| 1486 | 16 | 72 | P7 |

**Table S3:** Regions demonstrating change in MI over age. Low frequency and high frequency values refer to the frequency pair demonstrating the maximum t test value in a given cluster.

| Cluster size | Low Frequency (Hz) | High Frequency (Hz) | Channel |
| --- | --- | --- | --- |
| 15.60 | 6 | 44 | F8 |
| 19.10 | 16 | 84 | Fp1 |
| 27.99 | 12 | 88 | F4 |
| 34.46 | 12 | 52 | Fp1 |
| 41.95 | 20 | 40 | F7 |
| 51.40 | 20 | 84 | T8 |
| 52.02 | 16 | 84 | P7 |
| 71.67 | 12 | 88 | P8 |
| 105.3 | 12 | 80 | Pz |
| 203.3 | 12 | 60 | O1 |
| 313.1 | 14 | 72 | O2 |

**Table S4:** Regions demonstrating change in ϕ bias with age. Regions where ϕ bias decreased with age are indicated by negative cluster sizes, while regions where ϕ bias increased with age are indicated by positive cluster sizes. Low frequency and high frequency values refer to the frequency pair demonstrating the absolute maximum t test value.

| Cluster size | Low Frequency (Hz) | High Frequency (Hz) | Channel |
| --- | --- | --- | --- |
| -106.9 | 6 | 44 | F3 |
| -100.3 | 4 | 76 | Fp1 |
| -87.85 | 10 | 60 | T7 |
| -70.98 | 10 | 64 | F8 |
| -47.87 | 2 | 92 | Fz |
| -38.61 | 2 | 20 | F8 |
| -23.31 | 18 | 56 | F7 |
| 30.54 | 4 | 52 | O2 |
| 34.37 | 2 | 100 | C3 |
| 75.84 | 8 | 64 | P7 |
| 80.59 | 4 | 56 | Pz |
| 201.9 | 12 | 84 | Pz |
| 234.8 | 10 | 28 | P8 |
| 525.5 | 14 | 56 | O2 |
| 595.6 | 16 | 44 | O1 |

